# A moss N-Acetyltransferase-MAPK protein controls 2D to 3D developmental transition via acetylation and phosphorylation changes

**DOI:** 10.1101/2025.05.02.650421

**Authors:** Cloe de Luxán-Hernández, Thomas J. Ammitsøe, Jakob V. Kanne, Sabrina Stanimirovic, Milena E. Roux, Zoe Weeks, Michael Schutzbier, Gerhard Dürnberger, Elisabeth Roitinger, Liechi Zhang, Oliver Spadiut, Masaki Ishikawa, Mitsuyasu Hasebe, Laura A. Moody, Yasin F. Dagdas, Eleazar Rodriguez, Morten Petersen

**Author notes:** These authors contributed equally to this work (Cloe de Luxán-Hernández, Thomas J. Ammitsøe, Jakob V. Kanne).

## Abstract

Post-translational modifications (PTMs) finetune plant responses to developmental and environmental cues by impacting protein activity, stability, localization and interaction landscape. In this study we identified a moss specific protein which combines two common PTMs: acetylation and phosphorylation. This protein originated from the fusion of a MAPK with an N-acetyltransferase, for which we named it Rosetta NATD-MAPK 1 (RAK1). Using biochemical methods, we demonstrated that RAK1 has acetyltransferase activity that is enhanced by activation of its MAPK domain. Phenotypical studies of rak1 knockout mutants revealed a role for RAK1 in the regulation of the 2D-to-3D growth transition. Through Mass Spectrometry we verified that defective 2D-to-3D transition in the mutants was caused by differentially regulated acetylation and phosphorylation events associated to metabolic reprogramming and 3D differentiation. Collectively, this study uncovers a previously unknown multidomain protein and provides insights into the interplay of PTMs during developmental reprogramming.

**Teaser:** Acetylation and phosphorylation changes modulate the 2D to 3D developmental transition in *Physcomitrium patens*.

## Introduction

The shift from two-dimensional (2D) to three-dimensional (3D) growth was a key developmental event that facilitated the colonization of land by plants around 470 million years ago [1-3]. In recent years, diverse studies have focused on understanding the genetics behind this transition using the model bryophyte *Physcomitrium patens*. This is because the 2D phase of the *Physcomitrium patens* life cycle can be maintained indefinitely, and 3D growth is not essential for survival [4]. Thus, the transition from the 2D chloronema and caulonema filaments to the 3D gametophore buds can be easily observed. This makes *Physcomitrium patens* an ideal model for studying the molecular mechanisms involved in the shift from 2D to 3D growth in plants [1, 5]. Several key components involved in this developmental shift have been identified, including DEFECTIVE KERNEL 1 (DEK1), NO GAMETOPHORES 1 (NOG1), NO GAMETOPHORES 1 (NOG2), CLAVATA-like peptides and their conserved receptor RECEPTOR-LIKE PROTEIN KINASE 2 (RPK2) and the transcriptional co-repressor TOPLESS (TPL) family [6-10]. A particularly important role in this process is played by APETALA2-type (APB) transcription factors. Sustained *APB* expression in gametophore initial cells drives gametophore (3D) differentiation, whereas their downregulation promotes filamentous (2D) growth [11]. While significant progress has been made in understanding the transcriptional regulation of this transition even at the single cellular level [5, 9], little is known about how this process is controlled at the post-translational level. Post-translational modifications (PTMs) modulate different biological processes in plants such as signal transduction, stress responses, cell differentiation or metabolism by regulating the structure, function or localization of proteins [12, 13]. Some of the most investigated PTM in plants are protein phosphorylation and protein acetylation and they consist in the addition of a phosphoryl or an acetyl group to specific amino acid residues respectively [12]. In *Physcomitrium patens* specifically, protein phosphorylation has been linked to developmental reprogramming, immunity and abiotic stress responses [14-16], while protein acetylation has primarily been studied in the context of histone 3 (H3) during development, abiotic stress [17] and metabolism [18]. However, the roles of phosphorylation and acetylation in the 2D-to-3D growth transition remain unexplored. To address this gap, we identified a pair of homologous Rosetta proteins in *Physcomitrium patens*, which are fusion proteins that combine two or more distinct functional domains into a single polypeptide chain, enabling the study of complex molecular interactions [19-21]. These proteins in particular contain a full-length N-acetyltransferase class D (NATD) domain fused to a MAP kinase (MAPK) domain, leading us to designate them as Rosetta NATD-MAPK (RAK) 1 and 2. NATs catalyse the transfer of an acetyl group from acetyl-coenzyme A (Ac-CoA) to the N-terminus of specific substrates [22]. NATDs have been traditionally described to have a restricted substrate repertoire and are well known to acetylate the N-terminus of Histones H2A and H4 [23-26]. However, recent studies in yeast suggest additional targets including Transcriptional regulatory protein LGE1 [27]. Per contra, MAPKs are Serine/Threonine kinases that regulate adaptative and programmed responses by phosphorylating a wide range of substrates at conserved SP/TP dipeptide motifs [28]. Consequently, RAKs are remarkable in combining two post-translational modification enzymes which may regulate multiple cellular pathways, including the 2D to 3D growth transition. While the interplay between protein acetylation and phosphorylation is well documented in the modification of histone tails [29], a major mechanism in chromatin remodeling, direct links between them are few [30-32]. RAKs therefore represent a new tool for understanding protein modifications affecting chromatin, gene expression and cellular signaling. Our multiomics analyses revealed that loss of *RAK1* significantly impairs both the acetylome and phosphoproteome of *P. patens*. Moreover, we provide evidence of the developmental function of RAK1 in the induction of the 2D-to-3D transition by impacting proteins involved in metabolic reprogramming and previously identified regulators of this developmental process.

## Results

### *Physcomitrium patens* Rosetta NATD-MAPK (RAK) proteins arose by retrotransposition

The *Physcomitrium patens* genome encodes eight MAPKs clustered as four pairs [15]. We previously described two MAPKs involved in innate immunity [15], but the function of the remaining MAPKs has so far not been investigated. Interestingly, protein sequence analyses revealed that two MAPKs had a predicted size of ∼74 kD, which is significantly larger than the typical size of 40-45 kD observed in other MAPKs. This was explained by their abnormal structure: a MAPK preceded by a full-length N-acetyltransferase class D (NATD) (Fig. 1A). We named these proteins Rosetta NATD-MAPK 1 (RAK1) and -2 (RAK2). RAK1 (Pp3c9_11360) is encoded on chromosome 9 and shares approximately 82% protein identity with homologue RAK2 (Pp3c15_11610) that resides in a syntenic region on chromosome 15. The NATD sequences of RAK1 and RAK2 share six introns with a single homolog named NATD homolog (NATDH) (Pp3c17_14350) encoded on a non-syntenic region on chromosome 17. In contrast, the MAPK sequences of both proteins lack the 4-5 introns found in the other six moss MAPKs present on other chromosomes. The Rosetta therefore probably arose by retrotransposition of a MAPK C-terminally to a NATDH paralog, and this RAK was later duplicated on chromosome 9 or 15 (Fig. 1B). To better understand the origin of these Rosetta proteins, RAK1 paralogs were identified using OrthoFinder with the proteome of representative land plant, green algae and yeast species as input [33]. Phylogenetic trees generated using predicted NATD (Table S1) and MAPK sequences (Table S2) revealed that only the closely related moss *Ceratodon purpureus* has a homologous protein (CpRAK) containing both a NATD and a MAPK domain, while no other Rosetta proteins were found in other species (Fig S1A-B). High homology was still observed between predicted NATDs (Fig. S1A) and MAPKs (Fig. S1B) from other species and the corresponding *Physcomitrium patens* RAK domains, suggesting functional conservation (Fig. 1C and Fig S1A-B). Taken together, RAK proteins seem to be specific to mosses, while homologous MAPK and NATD proteins in other plants have remained separate.

**Figure 1.**
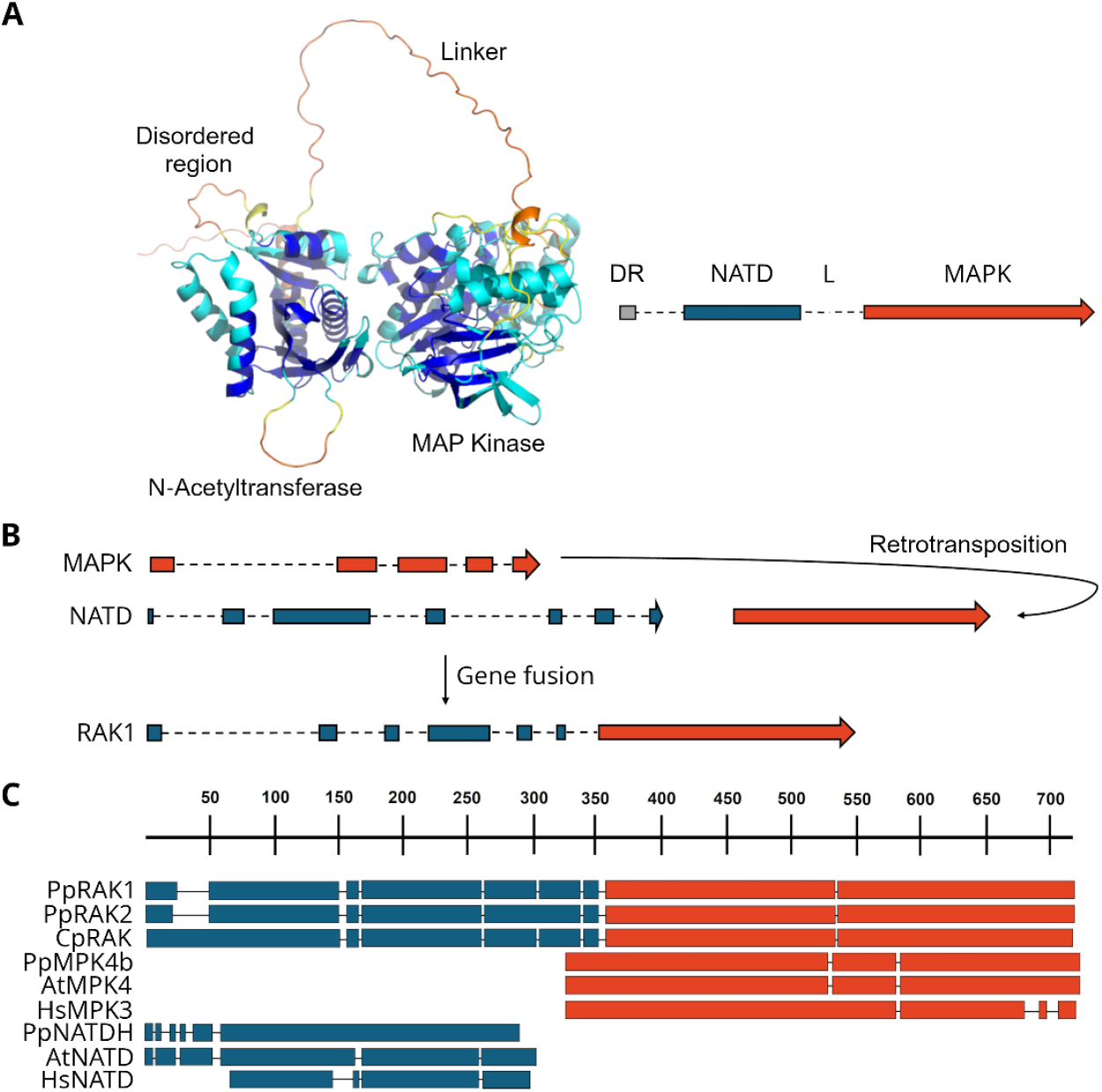
Rosetta NATD-MAPK proteins arose by C-terminal retrotransposition of a MAPK into an NATD and are specific to mosses. (A) Left panel: RAK1 protein structure model generated by Alphafold using default parameters Right panel: graphical simplified representation of the RAK1 fusion protein containing the disordered region (DR), N-Acetyltransferase (NATD) domain, linker region (L) and the MAP Kinase (MAPK) domain. (B) Graphic simplified representation of the C-terminal retrotransposition and subsequent gene fusion of an PpMAPK (orange) into an PpNATD (blue) to generate RAK1. Exons are represented as filled boxes and introns as dashed lines. PpMPK4b and PpNATDH were used as examples. (C) Multiple protein alignment using RAK1 as a reference made with the Constraint-based Multiple Alignment Tool (Cobalt) from NCBI based on conserved domain and local sequence similarity information. Protein sequences used for the alignment can be found in Table S3.

### RAK1 is important for the development of gametophores

To study the function of *RAK1* and *RAK2*, we attempted to generate KO lines for each *RAK* gene through homologous recombination, using a USER overhang-compatible system previously described in [15]. While for *RAK1* we were able to isolate several independent lines, the generation of KO mutants for *RAK2* was unsuccessful, suggesting that *RAK2* could be essential during protoplast recovery, which is the cell stage used for transformation during mutant generation. Therefore, further analyses were carried out only with *rak1* KO mutants (Fig. S2A and B). Phenotypical analyses revealed that the absence of *RAK1* leads to a reduction in the number of gametophores in 3 (Fig. 2A and B) and 4-week-old plants (Fig. S2C and D), accompanied by a slight decrease in gametophore height (Fig. 2C and D). No apparent protonema-derived defects were observed (Fig. 2A). Consequently, RAK1 seems to be specifically important during 3D growth.

**Figure 2.**
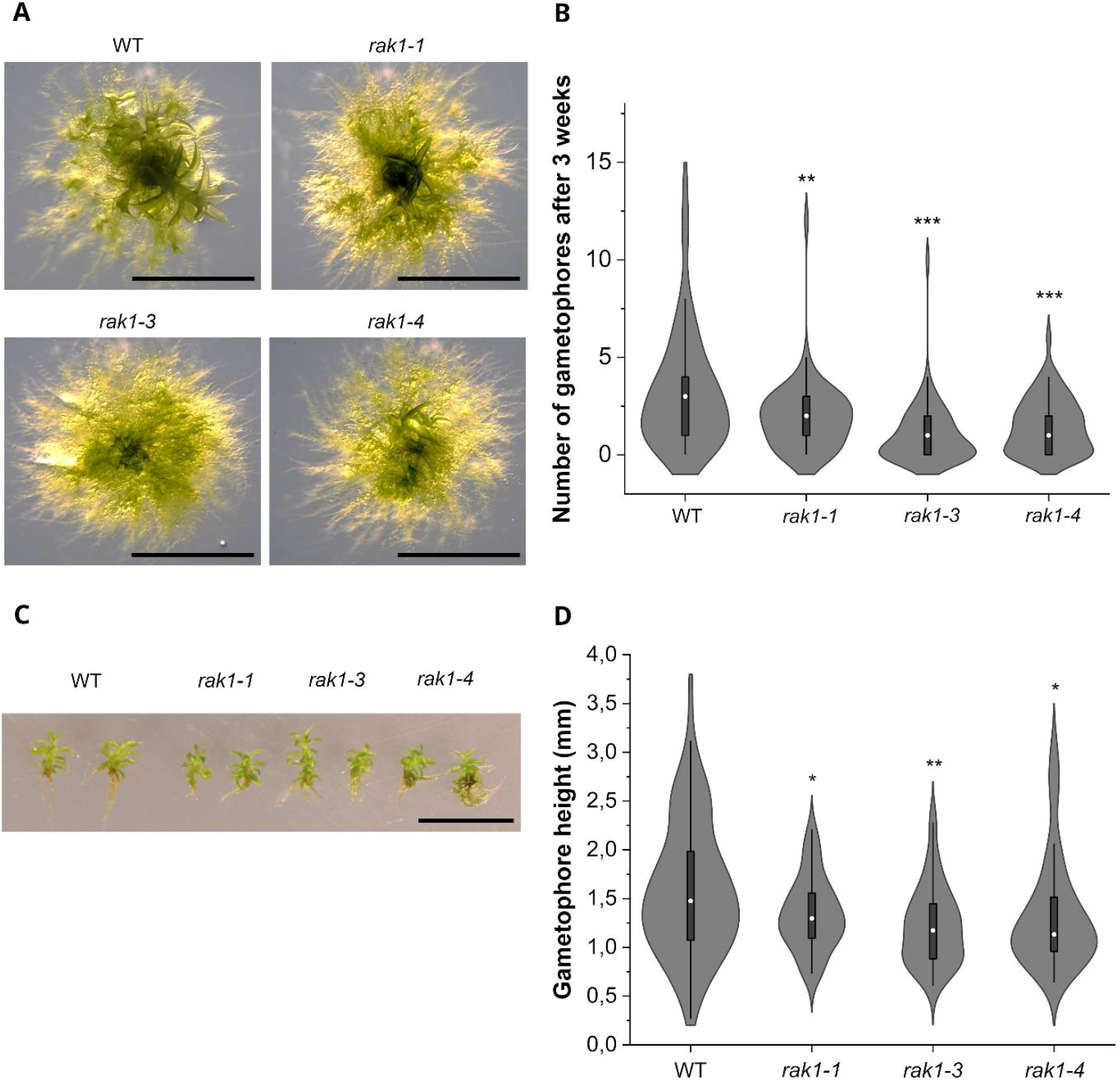
*rak1* mutants show decreased gametophore formation. (A) Representative phenotypes of 3-week-old W, *rak1-1, rak1-3* and *rak1-4* plants. Scale bar: 5 mm. (B) Quantification of the number of developed gametophores per plant: WT (N=58), *rak1-1* (N=61), *rak1-3* (N=64), and *rak1-4* (N=59). Statistical significance was calculated using a One-way ANOVA coupled to a Tukey test. Asterisks represent statistical difference against the WT: *rak1-1* (P value**: 0,00393), *rak1-3* (P value***: 5,69458×10-8), *rak1-4* (P value***: 3,54207×10-5). (C) Representative pictures of 4-week-old central gametophores in WT and *rak1* mutants. Scale bar: 1 cm. (D) Quantification of the height of 4-week-old single gametophores in WT (N=79), *rak1-1* (N=54), *rak1-3* (N=50), and *rak1-4* (N=70). Statistical significance was calculated using a One-way ANOVA coupled to a Tukey test. Asterisks represent statistical difference against the WT: *rak1-1* (P value*: 0,03696), *rak1-3* (P value**: 7,53847×10-4), *rak1-4* (P value*: 0,01355). Boxes represent the interquartile range 25th to 75th. Median shown as a white circle. Whiskers represent 1.5 IQR.

### RAK1 activity is required during bud initiation

Gametophores develop at side branch positions from precursor cells called gametophore initial cells [34, 35]. While more than 70% of side branches will acquire protonema identity, only approximately 2% become gametophore initial cells also known as buds [34] and this process is mediated by a crosstalk between auxin and cytokinin [35]. Furthermore, exogenous application of synthetic cytokinin benzylaminopurine (BAP) was shown to promote budding in mosses [11, 36]. Thus, to verify that the reduced number of gametophores was not caused by overall growth retardation in *rak1* mutants, the number of gametophore buds formed after BAP treatment was quantified (Fig. 3A and B). We observed a significant decrease in bud formation in *rak1* mutants, but these buds did not show any obvious developmental defects (Fig. 3A), which is consistent with what we observed in the fully developed gametophores (Fig. 2C). These findings suggest that RAK1 is important during bud initiation specification but not essential for correct bud development. Consistent with a role in bud formation, RAK1-GFP derived fluorescence (Fig. S2B) was observed in the nucleus and cytoplasm of gametophore buds (Fig. 3C). Additionally, the expression of *APB1-4* genes, a hallmark for bud formation [7] was also reduced in *rak1* mutants (Fig. 3D). Altogether, these data indicate that RAK1 is expressed in and is important for the specification of bud formation in gametophores.

**Figure 3.**
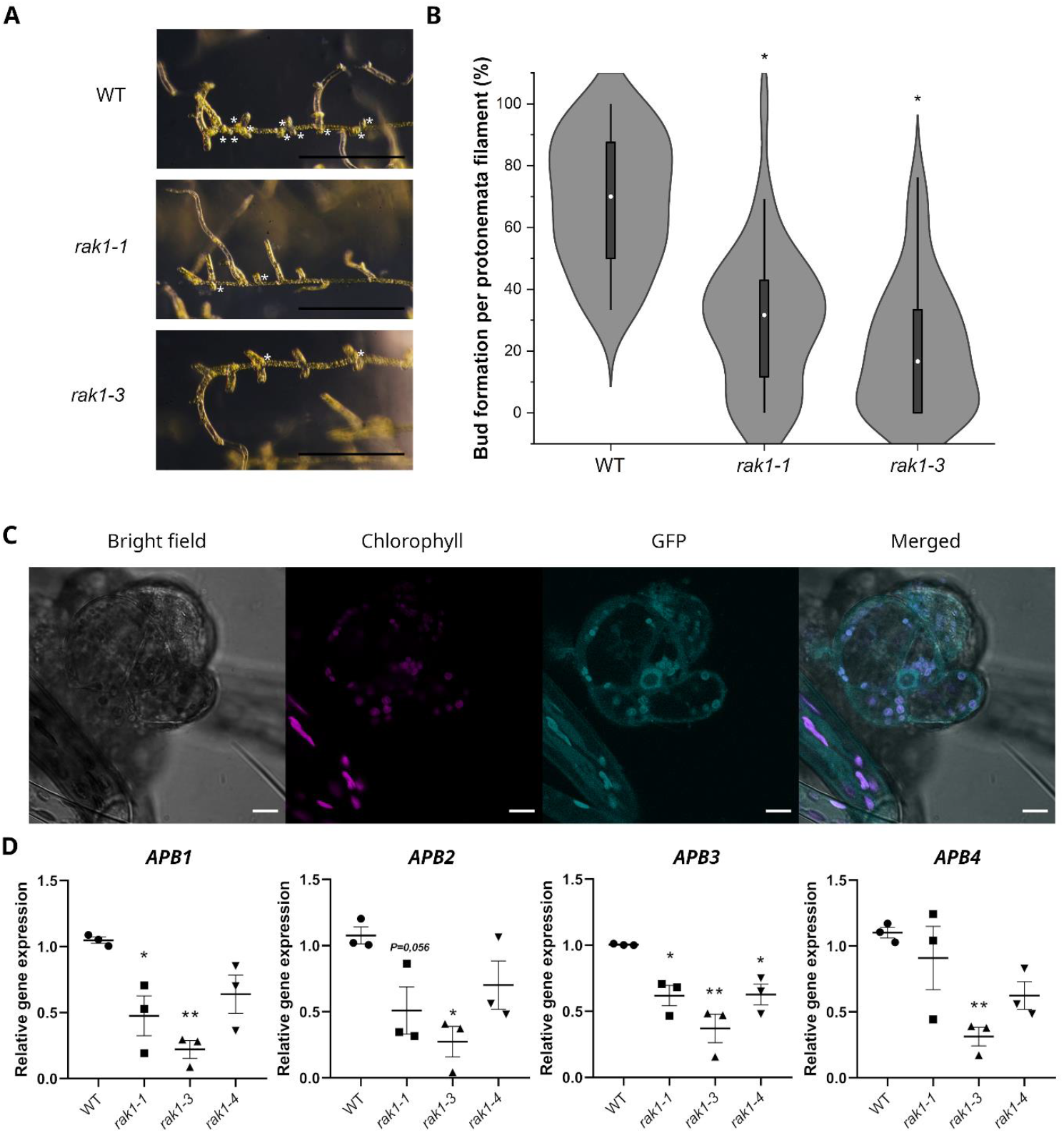
RAK1 is important to initiate bud formation. (A) Representative phenotypes of WT, *rak1-1* and *rak1-3* protonema filaments grown for 7 days in red light followed by growth in white light for 3 additional days after addition of BAP 1µM. Scale bar: 1 mm. White asterisks mark the position of buds. (B) Quantification of the percentage of buds per protonema filament in WT, *rak1-1* and *rak1-3* (N=40). The box represents the interquartile range 25th to 75th. Median shown as a white circle. Whiskers represent 1.5 IQR. Statistical significance was calculated using a One-way ANOVA coupled to a Tukey test. Asterisks represent statistical difference against the WT: (P value* <0,0001). (C) Representative confocal picture of RAK1-GFP derived fluorescence in the nucleus of a developing bud. Scale bar: 10 µm. (D) Relative expression of *APB1-4* genes in WT and *rak1* mutants. Average of 3 biological replicates with 3 technical replicates each ± SD is represented. Expression was normalized to *TUBULIN* and is relative to WT. Statistical significance was calculated using a One-way ANOVA coupled to a Dunnett test. Asterisks represent statistical significance against the WT in *APB1* expression: *rak1-1* (P value* 0.016) and *rak1-3* (P value** 0.0019); *APB2* expression: *rak1-3* (P value* 0.0107); *APB3* expression: *rak1-1* (P value* 0.0188), *rak1-3* (P value** 0.001) and *rak1-4* (P value* 0.021) and *APB4* expression: *rak1-3* (P value** 0.0093).

### Loss of RAK1 impacts both the acetylome and phosphoproteome of *Physcomitrium patens*

The unusual Rossetta structure of the *RAK1* gene with two post-translational modification domains, prompted us to ask whether the developmental defects observed in *rak1-1* mutants were caused by changes in protein levels. To that end, we analysed the proteome of 2-week-old plants of WT and *rak1-1* by mass spectrometry. Of the 9390 proteins that were detected in the analyses, 1447 and 1519 were upregulated and downregulated respectively in *rak1-1* in comparison to WT (Fig. 4A, Tables S4-5). Thus, loss of *RAK1* led to changes in 31.5% (2966) of the total detected proteome (9390). To gain more insight into the type of proteins altered in *rak1-1* we performed Gene Ontology (GO) term enrichment analysis. Translational and RNA processing processes were strongly enriched among the upregulated proteins (Fig. 4B, Table S6), whereas the downregulated proteins were associated to photosynthetic processes (Table S7, Fig. 4C). Despite the relatively large number of differentially expressed proteins in *rak1-1*, most of them showed only mild changes in their abundance in comparison to WT of less than two-fold changes (Fig. 4A). This indicates that RAK1 had a moderate impact on overall protein abundance, which might be insufficient to explain the phenotype of *rak1-1* mutants (Fig. 2A-D). However, PTMs have been shown to affect protein characteristics including interaction ability or localization [37]. Therefore, the defective 3D related traits in *rak1-1* mutants could be caused by changes in PTMs. To check this, we also analysed both the phosphoproteome (Fig. 4D-F) and global lysine acetylome (Fig. 4G and H) of 2-week-old WT and *rak1-1*. The phosphoproteomic analyses revealed that 1635 and 1317 proteins were significantly hyper- and hypophosphorylated respectively in *rak1-1* background (Tables S8-9). Since 480 proteins showed both hyper- and hypophosphorylation, a total of 2398 unique proteins had altered phosphorylation patterns in the absence of *RAK1*. Thus, 25,5% (2398) of the overall detected proteome (9390) is differentially phosphorylated in *rak1-1* mutants in comparison to WT (Fig. 4D). The almost equal distribution of hyper- and hypophosphorylated proteins suggests that most of these changes could be downstream effects derived from RAK1 activity, possibly through the regulation of transcription factors or kinases. Interestingly, GO term analyses showed enrichment of protein phosphorylation and gene expression related proteins within the differentially hyper and hypophosphorylated proteins in *rak1-1* (Fig. 4E-F and Table S10-11). Among them we found several AP2/ERF and BHLH putative transcription factors (Tables S8-9).

**Figure 4.**
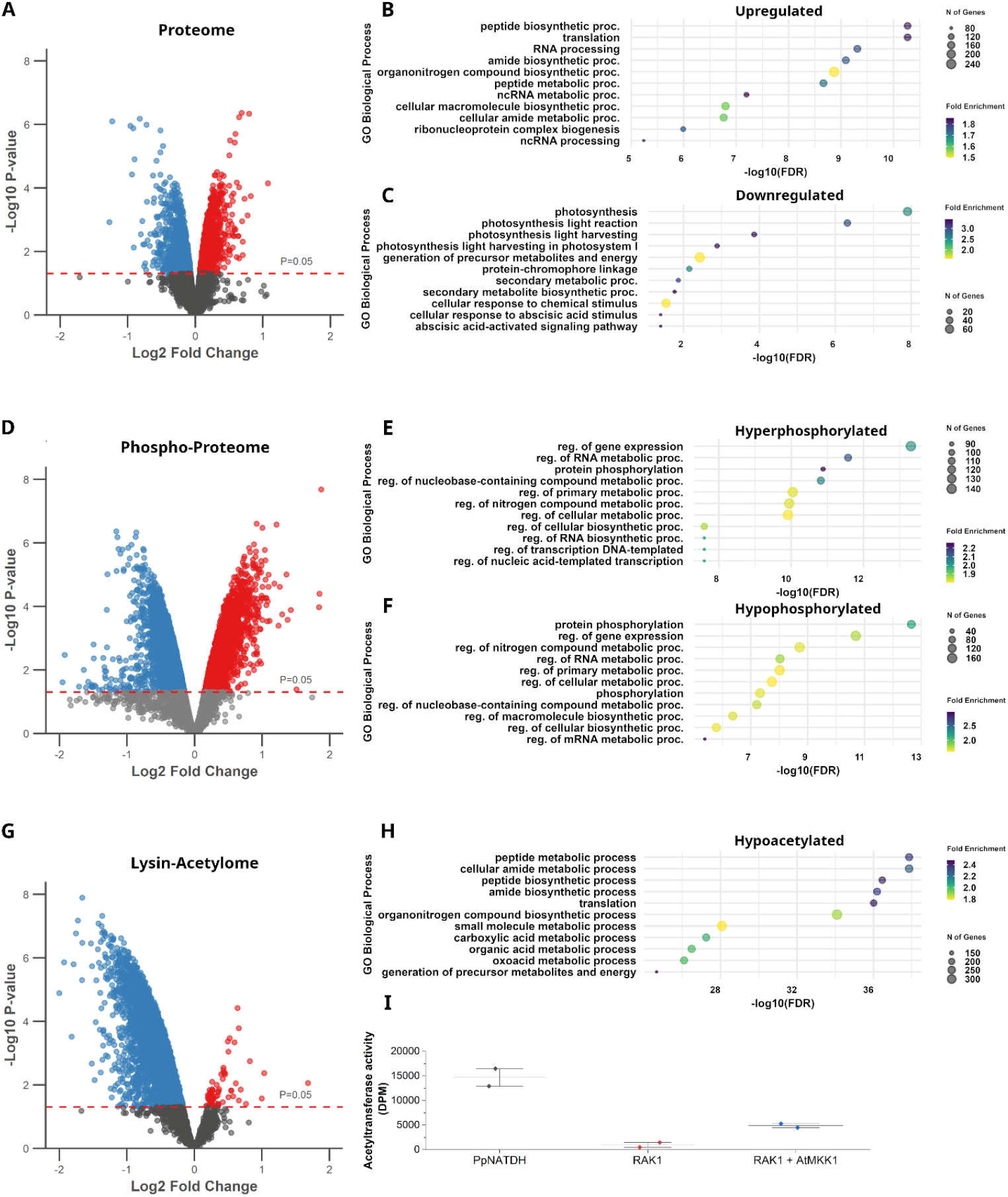
Absence of RAK1 leads to perturbed proteome, phospho-proteome and lysine acetylome. (A) Volcano plot representing the differentially upregulated (in red) and downregulated (in blue) proteins in *rak1-1* compared to WT with padj values < 0.05. (B-C) Representative top 11 significant Biological Process GO terms enriched in differentially upregulated (B) and downregulated (C) proteins identified in *rak1-1*. (D) Volcano plot representing the differentially hyperphosphorylated (in red) and hypophosphorylated (in blue) proteins in rak1-1 compared to WT with padj values < 0.05. (E-F) Representative top 11 significant Biological Process GO terms enriched in differentially hyperphosphorylated (E) and hypophosphorylated (F) proteins identified in *rak1-1*. (G) Volcano plot representing the differentially hyperacetylated (in red) and hypoacetylated (in blue) proteins in *rak1-1* compared to WT with padj values < 0.05. (H) Representative top 11 significant Biological Process GO terms enriched in differentially hypoacetylated proteins identified in *rak1-1*. (I) In vitro acetyl transferase activity showing incorporation of radioactive acetyl-CoA onto a H4 peptide after addition of PpNATDH, RAK1 or RAK1 together with AtMKK1. Two technical replicates and their average is represented.

These findings suggest that the altered phosphoproteome in *rak1-1* may influence transcriptional regulation and potentially contribute to some of the observed proteomic changes in *rak1-1* (Fig. 4A). Additionally, 105 potential kinases predicted to be involved in different biological processes were associated to the GO term “Protein Phosphorylation” (Table S11). These include PHOTA1/A2/B1/B2 (Q6BCU1, Q6BCU0, Q6BCT8, Q6BCT7), involved in photomorphogenesis; ATG13 (A9S9Z6) with a role in autophagy; the DNA damage repair protein ATM (A0A2K1L2S2) and energy monitoring proteins SNF1a/b (Q6V8Y3, Q6V8Y5) and RAPTOR (A0A2K1IE61). Thus, changes in phosphorylation status of these kinases might disturb a myriad of biological processes, further contributing to the proteomic changes found in *rak1-1*. Since RAK1 is predicted to contain a MAPK, to gain a deeper insight into potential direct phosphorylation targets, we searched for peptides hypophosphorylated at SP or TP dipeptide motifs and found 934 potential targets (Table S12). Among those with the highest Log2fold changes was a Tudor domain-containing protein (A0A2K1J4F2). Tudor motif containing proteins were previously described in humans to facilitate the recruitment of POLYCOMB REPRESSIVE COMPLEX 2 (PRC2) by recognizing methylated lysines and consequently acting as histone readers [38]. Therefore, RAK1 could also directly affect the epigenetic landscape of *Physcomitrium patens*, which could lead to additional transcriptional changes. In this context, since NATDs are known to regulate epigenetic changes by specifically acetylating H4, we also tested the impact of RAK1 on the global lysine acetylome. 1733 unique proteins were found to be significantly hypoacetylated in *rak1-1*, whereas only 61 were hyperacetylated (Fig. 4G, Tables S13-14). This means that 19.1% (1733) of the detected proteome (9390) had significantly perturbed acetylation patterns. Contrary to the proteome and phosphoproteome analyses, lysine acetylation showed a strong tendency towards hypoacetylation in the absence of *RAK1*. GO term analysis revealed that hypoacetylated proteins in *rak1-1* were associated with translation, including ribosomal subunits and metabolism related processes (Fig. 4H, Table S15). This aligns with studies linking metabolic changes to gametophore formation [39] and the broader role of protein acetylation in metabolic regulation [18, 40, 41]. Interestingly, among the enriched hypoacetylated proteins, we identified key enzymes involved in pyruvate synthesis (PIRUVATE KINASE) and its conversion into Acetyl-CoA (PYRUVATE DEHYDROGENASE E1 and DIHYDROLIPOAMIDE ACETYLTRANSFERASE E2) (Fig. S3, Table S16). Given that NATD homologs in human and yeast regulate cellular Acetyl-CoA levels and glycolysis gene expression [42, 43], these findings suggest that RAK1 may influence Acetyl-CoA production. Additionally, Histone H4, a known target of NATDs in other organisms, was found to be hypoacetylated in 5 positions in *rak1-1*: 3 in the tail (H4K101ac, H4K105ac and H4K109ac), corresponding to conserved yeast and human acetylation sites (H4K8ac, H4K12ac, H4K16ac) [44-46] and 2 in the histone fold (H4K172ac and H4K184ac) (Table S13). Despite H4 hypoacetylation at multiple residues in *rak1-1* mutants, it cannot be excluded as a downstream effect resulting from the absence of *RAK1*. To further prove whether RAK1 could directly acetylate H4, we conducted an in vitro acetylation assay combining the 19-mer highly conserved H4 tail (Fig. 4I) with RAK1 and PpNATDH in the presence of [14C]-Acetyl-CoA. Acetyl group incorporation on the H4 peptide was observed in the presence of RAK1 (Fig. 4I), further supporting H4 as a target of RAK1. However, the signal was weaker compared to that generated by PpNATDH. The apparent lower histone acetyltransferase (HAT) activity of RAK1 could be caused by the C-terminal retrotransposition of a MAPK onto its NATD domain. Since the NATD domain has two potential phosphorylation sites for MAPKs, we wondered whether the HAT activity is dependent on MAPK activation. Based on MAPKs conserved activation by MAP KINASE KINASES (MKKs), a constitutively active AtMKK1 was added together with RAK1 and H4 to induce the activation of the MAPK domain in RAK1 (Fig. 4I). This resulted in a marked increase in H4 acetylation, indicating that RAK1 HAT activity is dependent on auto-MAPK activity. Therefore, the RAK1 NATD domain could be a direct target of the MAPK domain through autophosphorylation.

### RAK1 influences the phosphorylation and abundance of proteins involved in the 2D to 3D growth transition

Our data showed that RAK1 can regulate the acetylation status of H4, which could subsequently regulate the expression and activity of downstream genes involved in different biological processes. To understand the role of RAK1 in the context of the 2D to 3D developmental transition, we assessed the abundance, phosphorylation and lysine acetylation status of key proteins involved in this change in *rak1-1* mutants (Fig. 5). Surprisingly, despite the large number of *rak1* hypoacetylated proteins, none of the targets found in the literature was regulated by acetylation. However, several proteins were regulated in either their phosphorylation status or protein abundance (Fig. 5, Table S17). Among the differentially phosphorylated proteins we found NOG1, FLOE2L-1, CRL1/2, AN1-1 and TPL1/2. Notably, we observed that both the essential 2D-to-3D transition protein NOG1 [7] and its suppressor FLOE2L-1, which acts as a negative regulator of 3D growth initiation [47], were hypophosphorylated in *rak1-1*. Interestingly, FLOE2L-1 displayed the highest level of hypophosphorylation among the candidate proteins (Fig. 5, Table S17). Regarding the up or downregulated proteins, only mild differences were observed between *rak1-1* and WT. We found the 2D-3D growth involved PRC2 protein, CLF [48], to be downregulated along with CRL2, ETR1/2B/3, SNF1b, TEL1 and VNS5. Surprisingly, we found NOG2 to be upregulated in *rak1-1* together with the transcriptional co-repressors TPL1 and TPL2 [9]. Taken together, our data shows that known gametophore development proteins are regulated via phosphorylation and to some extent protein abundance but not through lysine acetylation.

**Figure 5.**
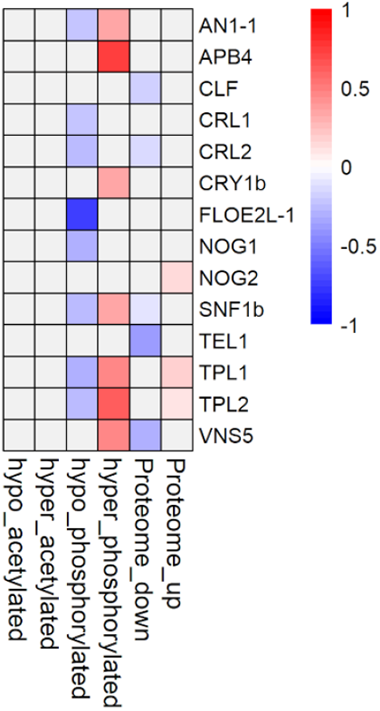
Proteins involved in the 2D to 3D growth transition are differentially regulated in *rak1-1*. Heat map showing differential Log2Fold changes in *rak1-1* mutants in lysine acetylation, protein phosphorylation and protein abundance of known proteins involved in the 2D-to-3D transition in *P. patens*.

### *rak1* mutants exhibit traits linked to enhanced protonema identity

The reduced gametophore development observed in *rak1* mutants (Fig. 2A and B), combined with the altered phosphorylation status and abundance of several proteins involved in the 2D-to-3D transition (Fig. 5), led us to hypothesize that protonema growth might be enhanced in the absence of *RAK1*. To this end, the surface area of 3-week-old plants was quantified (Fig. 6A). *rak1* mutants showed an increase in overall protonema size (Fig. 6A), suggesting that their defective transition into 3D growth might lead to 2D expansion. Therefore, to further investigate whether *rak1* mutants’ 2D and 3D identities were impaired, the expression of gametophore and protonema marker genes was evaluated (Fig. 6B). *RIBULOSE-BIPHOSPHATE CARBOXILASE* (*RuBP*), a protonema marker [49], was expressed similarly in WT and *rak1* mutants, whereas the gametophore marker [49] *FASCICLIN-LIKE ARABINOGALACTAN PROTEIN 11* (*FASC*) was significantly less transcribed in the absence of *RAK1* (Fig 6B). This indicates that *rak1* mutants might have a reduced gametophore identity. To further investigate this hypothesis, we compared our proteomic data (Fig. 4A) with previously published [50] protonema and gametophore-enriched proteins (Fig. 6C, Table S18). Protonema enriched proteins were statistically overrepresented in upregulated proteins in *rak1-1* background, while gametophore enriched proteins were only overrepresented among the downregulated (Fig. 6C). Additionally, a comparison with previously published gametophore acetylome data [18] revealed that 52% (326/624) of the gametophore-specific acetylated proteins overlapped with hypoacetylated proteins in rak1-1 (Fig. 6D, Table S19). Interestingly, GO term enrichment analyses revealed that these overlapping proteins are involved in metabolic processes (Fig. S4, Table S20). Taken together, these findings suggest that cell fate specification is impaired in *rak1* mutants and supports a role for RAK1 in the PTM regulation of gametophore initiation.

**Figure 6.**
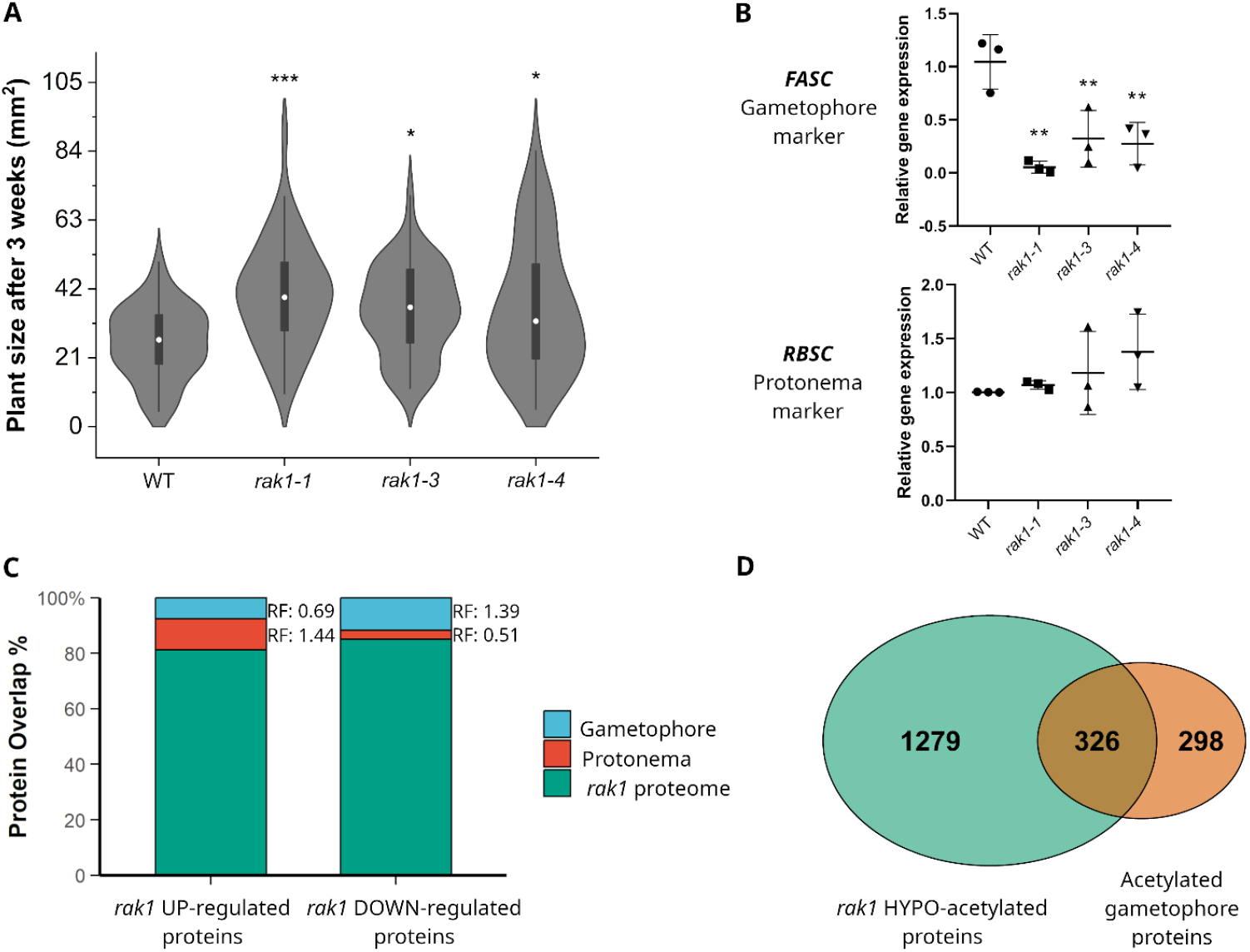
*rak1* mutants show traits linked to impaired protonema and gametophore identity. (A) Plant size of 3-week-old WT (N= 50), *rak1-1* (N=49), *rak1-3* (N=51) and *rak1-4* (N=50). The box represents the interquartile range 25th to 75th. Median shown as a white circle. Whiskers represent 1.5 IQR. Statistical significance was calculated using a One-way ANOVA coupled to a Tukey test. Asterisks represent statistical difference against the WT: *rak1-1* (P value*** 2,45676×10-5), *rak1-3* (P value* 0,00764) and *rak1-4* (P value* 0,00564). (B) Relative expression of protonema marker *RBSC* (Pp3c2_4100) and gametophore marker *FASC* (Pp3c26_11940) in 2-week-old plants of WT and *rak1* mutants. Average of 3 biological replicates with 3 technical replicates each ± SD is represented. Expression was normalized to *TUBULIN* and is relative to WT. Statistical significance was calculated using a One-way ANOVA coupled to a Dunnett test. Asterisks represent statistical significance in *FASC* expression against the WT: *rak1-1* (P value** 0,0011), *rak1-3* (P value** 0,0079) and *rak1-4* (P value** 0,0055). (C) Graphical representation of the overlap between gametophore (blue) or protonema (red) enriched proteins found in [50] and the differentially upregulated (left) and downregulated (right) *rak1* proteins (green). Statistical significance for each overlap was calculated using hypergeometric tests. Representation factors (RF) >1 indicate overrepresentation. (RF:0.69/P value:7,3629×10-7, RF:1.44/P value:1,3294×10-7, RF:1.39/P value:8,2096×10-8, RF:0.51/P value:1,4243×10-8). (D) Venn diagram representing the overlap between *rak1* differentially hypoacetylated proteins (green) and the gametophore acetylated proteins found in [18].

## Discussion

In this paper we identified a Rosetta NATD-MAPK protein that combines two of the most common PTMs, which are acetylation and phosphorylation. *Phylogenetic analyses* revealed that RAK proteins are only present in the closely related mosses Physcomitrium patens and *Ceratodon purpureus*. No homologous sequences containing both a NATD and a MAPK were identified in any other plant lineage (Fig. S1A and B). Thus, the origin of RAK proteins likely occurred by the retrotransposition of a MAPK into a NATD in their common ancestor between 368 and 160 MYA, the estimated diverging time of Sphagnum and Ceratodon respectively [51].

Through a combination of biochemical and mass spectrometry assays (Fig. 4G-I), we show that RAK1 directly acetylates H4 possibly functioning as a NATD. The global acetylome in *rak1* mutants additionally revealed how H4K8, H4K12 and H4K16 acetylation was dependent on RAK1. Interestingly, these acetylation sites were previously described in other organisms such as Arabidopsis, yeast and human H4 [44-46, 52] and likely act as regulatory checkpoints [52]. Additionally, NATD N-terminal H4 acetylation has been suggested to enable local, but not global, H4K5ac acetylation to promote the expression of transcription factors [32]. Thus, RAK1 mediated H4 N-terminal acetylation could promote subsequent acetylation of H4 by HATs. Surprisingly, we also observed that RAK1s acetyltransferase activity is regulated by the activation of their MAPK domain (Fig 4I). Consequently, this suggests that RAK1 regulates itself and both domains have been kept together to ensure rapid activation of the NATD domain, which aligns with the Rosetta Stone Protein hypothesis [19, 53].

Given the importance of acetylation and phosphorylation to protein function, it was expected that loss of function of RAK1 would lead to phenotypical consequences. Precisely, we observed that *rak1* mutants show defects in the transition from 2D to 3D development, namely due to reduced bud specification (Fig. 2 and Fig. 3). Bud specification requires cytokinin-induced reprograming of somatic cells to become multiplanar stem cells [54, 55] which eventually differentiate into gametophores [2, 35]. As observed in multicellular eukaryotes, reprogramming requires epigenetic, transcriptional, metabolic and proteome remodelling [56-58]. In this context, our acetylome data provides evidence of defective proteome in *rak1* mutants which should interfere with reprogramming. For instance, H4 residues detected in our acetylome experiment have been associated with transcriptional upregulation via nucleosome relaxation and enhanced gene accessibility [59]. Thus, it should be expected that hypoacetylation of H4 would lead to downregulation of several genes and indeed we observed downregulation of the bud-defining TF *APB*s (Fig. 3D). Further support for defective reprogramming comes from the fact that there is an enrichment for hypoacetylation of metabolic processes like pyruvate-to-Acetyl-CoA conversion and TCA cycle (Fig. 4H and Fig. S3). These findings are consistent with previous reports showing that TCA cycle enzymes can be modulated by acetylation in *Physcomitrium patens* [18]. Moreover, NATD homologs are known to regulate acetyl-CoA levels via a yet undetermined mechanism [42, 43] and directly associate with ribosomes to acetylate nascent proteins [60]. So, it is plausible that RAK1 direct modulation of these metabolic enzymes affects the pool of Acetyl-CoA and this both affects the metabolic and epigenetic reprogramming necessary to kickstart bud formation. In support of this, a body of evidence has been found linking metabolic reprogramming as inducers of developmental processes such as fate specification, differentiation and stem cell maintenance in different organisms [61-64].

Further supporting a role for RAK1 during developmental reprogramming, our proteomic analysis in *rak1-1* mutants revealed that important regulators of the 2D-to-3D transition showed changes in protein abundance and phosphorylation status (Fig. 5). Among the hypophosphorylated proteins, we found 5 containing the typical SP or TP dipeptide phosphorylation site of MAPKs (Tables S12,17). Therefore, it cannot be discarded that besides phosphorylating itself, RAK1 might directly phosphorylate some of these 2D-to-3D regulators. Among those we find FLOE2L-1 [47]. Mutation of *FLOE2L-1* in *nog1* mutant background was described to recover the cell division orientation defects associated to the *NOG1* disruption in most cases, while only partially restoring the formation of gametophore initial cells [47]. Interestingly, *rak1* mutants also showed impaired formation of gametophore initial cells after BAP treatment (Fig 3A and Fig. 3B), while the formed buds were WT-like (Fig. 3A). Based on this, it can be suggested that RAK1 dependent phosphorylation of FLOE2L-1 may impact its stability or activity. In the absence of RAK1, FLOE2L-1’s predicted ability to degrade a 2D-to-3D repressor [47] would be impaired, leading to the reduced gametophores seen in *rak1* mutants (Fig. 2A and B). Supporting this, the closest homolog of FLOE2L-1 in *Arabidopsis thaliana*, AtFLOE1, was shown to be phosphorylated by the TOR kinase [65], indicating that AtFLOE1’s protein stability might also depend on its phosphorylation. In conclusion, we have identified a yet uncharacterized Rosetta protein, comprising the fusion of NATD and MAP Kinase domains, which is an important regulator of 2D to 3D development in mosses. While our data suggest that RAK1 is essential for maintaining acetylome and phosphoproteome homeostasis to enable 3D growth, future efforts should focus on identifying the direct targets of the NATD and MAPK domains of RAK1, as well as further investigating their interplay.

## Materials and Methods

### *Physcomitrium patens* growth conditions

*Physcomitrium patens* (Gransden 2004 strain) was grown at 21°C 55µEm^-2^s^-1^, with a 16 h light /8 h dark cycle on cellophane-overlaid 92 mm Sarstedt BCD-AT media plates (MgSO_4_.7H_2_O 250 mg/L, KH_2_PO_4_ 250 mg/L, KNO_3_ 1010 mg/L, Ammonium tartrate 920 mg/L, FeSO_4_.7H_2_O 12.5 mg/L, CaCl_2_.H_2_O 147 mg/L, trace elements 1 mL/L [H_3_BO_3_ 61mg/l, (NH_4_)_6_Mo_7_O_24_·4H_2_O 38mg/L, CuSO_4_·5H_2_O 6mg/L, CoCl_2_·6H_2_O 5,1mg/L, ZnCl_2_ 4,1mg/L, MnCl_2_·4H_2_O 4,1mg/L], pH 6.5 adjusted with KOH, agar 8g/L.). For protoplast regeneration, BCDAT media was supplemented with 6% D-mannitol and 5mL/L of CaCl_2_ 1M.

### Phenotypical analyses

Gametophore formation was assessed by following growth upon reprogramming of single excised 3-week-old gametophore leaves for up to 4 weeks on BCDAT media plates without cellophane. The number of gametophores in each plant was counted using a Leica M165 FC Fluorescent Stereo Microscope coupled to a Sony alpha 6000 camera. Individual plates were also scanned with a Ricoh IM C5500 printer to subsequently measure plant area for each timepoint using ImageJ.

Counting of newly formed gametophore buds was performed by growing small pieces of 7-day-old protonema cultured on BCDAT medium on 35 mm Nunclon plates filled with 4 mL of solid BCDAT. 300 μL of sterile water were added on top of the tissue and covered with a cover glass. Subsequently plates were placed in dark boxes with a red-light filter under continuous light at 25ºC. After 7 days, the plates were taken out of the black boxes and 300 μL of BAP 1 μM were added. After 72 h of BAP addition, bud formation was calculated by dividing the number of buds between the total number of buds and side branches on 40 protonema filaments per line.

### Phylogenetic analyses

Proteome sequences (primary transcript only) were obtained for the following species; *Arabidopsis thaliana* Araport11 [66], *Saccharomyces cerevisiae* R64-1-1 [67, 68], *Solanum lycopersicum* ITAG2.4 [69], *Populus trichocarpa* v3.1 [70], *Chara braunii* S276v1.0 [71], *Chlamydomonas reinhardtii* v5.5 [72], *Anthoceros agrestis* Oxford [73], *Azolla filiculoides* v1.1 [74], *Gingko biloba* v2021 [75], *Glycine max* Wm82 ISU-01 v2.1 (DOE-JGI, http://phytozome.jgi.doe.gov), *Gossypium raimondii* v2.1 [76], *Musa acuminata* v1 [77], *Panicum hallii* v3.2 [78], *Salvinia cucullata* v1.2 [74], *Thuja plicata* v3.1 [79], *Physcomitrium patens* v3.3 [80], *Marchantia polymorpha* v3.1 [81], *Selaginella moellendorffii* v1.0 [82], *Oryza sativa* v7.0 [83], *Zea mays* PH207 v1.1[84], *Sorghum bicolor* v3.1.1 [85], *Brachypodium distachyon* v3.1 [86], *Ceratodon purpureus* R40 v1.1 [87], *Ceratopteris richardii* v2.1 [88], *Chlorokybus atmophyticus* CCAC 0220 v1.1 [89], *Klebsormidium nitens* NIES-2285 v1.1 [90], *Spirogloea muscicola* CCAC 0214 [91], *Mesotaenium endlicherianum* SAG 12.97 [91], *Volvox carteri* v2.1 [92], *Sphagnum fallax* v1.1 and *Sphagnum magellanicum* [93], *Diphasiastrum complanatum* v3.1 (DOE-JGI, http://phytozome-next.jgi.doe.gov/).

RAK paralogues were identified by running OrthoFinder on the proteome data [33]. MAFFT was used to align the sequences. IQ-TREE with automatic model selection and ultrafast bootstrapping was used to generate the gene tree [94-97].

The multiple alignment for schematic visualization of the conservation of the NATD and MAPK domains in selected taxa (Table S3) was made with the NCBI Constraint-based Multiple Alignment Tool (Cobalt) using default parameters. The multiple alignment for schematic visualization of the conservation of the NATD and MAPK domains in selected taxa (Table S3) was made with the NCBI Constraint-based Multiple Alignment Tool (Cobalt) using default parameters.

### Generation of mutant lines

Knockout mutants for *RAK1* were generated via homologous recombination following the protocol described by [15]. In brief, approximately 1000 base pairs (bp) upstream (Left Border) and downstream (Right Border) of the *RAK1* genomic sequence were amplified using primers with USER-compatible overhangs. The resulting PCR products were inserted into the USER-cloning vector pMBLU, which had been digested with PacI and AsiSI and nicked with Nt.BbvCI. For the RAK1-GFP construct, the same Right Border was used, while the Left Border was replaced with a sequence spanning the 3’end of the genomic region of *RAK1* but excluding the STOP codon. This construct was cloned using the same USER-cloning strategy into the pMBLU-GFP vector. Subsequently, 30 μg of linearized vectors were transformed in *Physcomitrium patens* protoplasts using polyethylene glycol following the protocol described by [98]. Stable transformants were selected by culturing protoplasts on selective media (50 mg/mL G418) for 2 weeks, followed by 2 weeks on non-selective media, and a final 2 weeks on selective media. The generation of the 35S::GFP line used during Western Blot analyses was described in [15].

### Western Blot analyses of the *rak1* KO mutants and the RAK1-GFP KI line

Total protein extracts were extracted from 2-week-old plants as previously described in [15] using Lacus buffer (50 mM Tris-HCl, pH7.5, 10mM MgCl2, 15mM EGTA, 100mM NaCl, 2mM DTT, 30mM b-glycerol phosphate, and 0.1% Nonidet P-40) supplemented with phosphatase inhibitor (PhosSTOP; Roche) and protease inhibitor cocktails (Complete; Roche). Samples were subsequently cleared by centrifugation at 4ºC for 30 min and boiled for 5 min at 95ºC in SDS Loading buffer. Total protein was separated using 12% SDS-PAGE gels (Biorad) and electroblotted for 3 hours at 120 V. After wet transfer ON at 4ºC, immunoblots were blocked for 1 hour in TBS-Tween (0,1% v/v) and 5% BSA. Phosphorylation of native RAK1 and RAK-GFP was detected by incubating with anti TeY antibody (1/2000; Cell signaling) for 2 hours at room temperature. This was followed by incubation with an anti-rabbit IgG HRP (1/5000; Promega) for 1 hour at room temperature. The horseradish peroxidase-conjugated antibody was visualized with ECL substrate (2mM 41BPA, 500 µM luminol, 100mM Tris pH 8,8 and 1,7×10-2 H2O2) using a Sony A7S camera. The membrane was subsequently incubated ON at 4ºC with an anti-GFP antibody (1/5000; Proteintech), followed by incubation for 1 hour at room temperature with an anti-mouse AP (1/5000; Promega). Detection was done by incubating in NBT/BCIP Stock solution (Roche) following manufacturer’s instructions. Membrane was pictured using a Canon EOS 1200D camera.

### Imaging

Representative pictures of 3-and 4-week-old plants were taken using a Leica M165 FC Fluorescent Stereo Microscope coupled to a Sony alpha 6000 camera. Individual gametophores after 4 weeks were additionally photographed using a Canon EOS 1200D camera.

Confocal images of RAK1 derived GFP fluorescence were obtained using an LSM-700 Zeiss confocal microscope. Images were processed with the Zen Blue 2012 software.

### Transcriptional analyses

Total RNA from 2-week-old plants was extracted using the NucleoSpin RNA kit (Macherey Nagel) according to manufacturer’s instructions. 1 µg of newly isolated RNA was afterwards treated with DNAse I (Thermo Fisher Scientific) and reverse transcribed into cDNA using the RevertAid First Strand cDNA Synthesis Kit (Thermo Fisher Scientific) following manufacturer’s instructions. SYBR Green Master Mix (Thermo Fisher Scientifc) and the QuantStudio 5 cycler (Thermo Fisher Scientific) were used for performing the quantitative PCRs. Generated data were quality-controlled and normalized to the reference gene *TUBULIN* using the QuantStudio Design and Analysis software. All experiments were repeated at least three times each in technical triplicates.

### Filter-aided sample preparation (FASP) for Mass spectrometry

2-week-old tissue grown on BCDAT plates overlaid with cellophane was generated in triplicates as tissue source. Samples were processed using the FASP method with minor modifications [99]. Briefly the 1mg of plant powder was dissolved in 1500 µl of SDT buffer (4% SDS, 0.1M DTT in 0.1M Tris pH=7.5) and incubated 10 min at 95°C. Then samples were sonicated with ultrasonication probe connected to Ultrasonic processor UP100H (Hielscher) with 30 pulses (0.5s, 50% amplitude). The cell debris was pelleted at 14000 × g for 15 min. The total protein concentration was determined using the 660nm Assay (Thermo Scientific). Plant cell lysate (approx. 1.2 mg Protein) was mixed with 9 ml of UA solution (8M urea in 0.1M TRIS; pH=8.5) Amicon Ultra-15 centrifugal filter unit with a 30 kDa molecular weight cutoff (Millipore). All buffer exchanges were performed by centrifugation at 14000 × g at room temperature. The first buffer exchange (UA buffer) was repeated, followed by alkylation of proteins by addition of 200 μl of 0.1M iodoacetamide in UA for 30 min at room temperature. After the alkylation two buffer exchanges using 8M UA buffer, followed by two further buffer changes using 0.8 M urea, 50 mm Triethylammonium bicarbonate (TEAB, Fluka). For the Trypsin digestion 800 μl of 50mM TEAB Buffer containing Trypsin (Trypsin Gold, Promega, Madison, WI) at an enzyme to protein ratio of 1:100 was added and incubated at 37 °C overnight. Generated peptides were collected by centrifugation. This was followed by 2 washes of 500 μl 50mM TEAB. About 0.6mg (in 200µl 100mM Hepes pH= 7.6) were labelled with one separate channel of the TMT 6 plex reagent (Thermo Fisher)/TMT6plex) according to the manufacturer’s description. The labelling efficiency was determined by LC-MS/MS on a small aliquot of each sample. Samples were mixed in equimolar amounts and equimolarity was again evaluated by LC-MS/MS. The mixed sample was acidified to a pH below 2 with 10% TFA and was desalted using C18 cartridges (Sep-Pak Vac 1cc (200mg), Waters). Peptides were eluted with 2 × 450 µl 80% Acetonitrile (ACN) and 0.1% Formic Acid (FA), followed by freeze-drying. Subsequently, neutral pH fractionation was performed as described in [100], with minor modifications. A 30 min gradient of 4.5% to 45% ACN (VWR, 83639.320) in 10 mM ammonium formate (1 ml formic acid (26N) (Merck), 3 ml ammonia (13N)in 300 ml H2O, pH = 7 - 8, dilute 1:10) was used on an Vanquish FLEX HPLC System (Dionex, Thermo Fisher Scientific) equipped with a XBridge Peptide BEH C18 (130 Å, 3.5 μm, 4.6 mm x 250 mm) column (Waters) (flow rate of 1.0 ml/min). 40 fractions were collected and subsequently pooled in a non-contiguous manner into 20 pools and quantified using a monolithic HPLC system. An aliquot of 5 µg of peptides was taken from each of these fractions for the proteome measurements. The 20 Fractions were pooled again into 5 pools and a C18 cartridge (Sep-Pak Vac 1cc (50mg), Waters). The organic content of the eluates was removed by using the speed vac and lyophilized overnight.

### Acetylated peptide enrichment

Acetylated peptides were enriched, washed, eluted and desalted as described in [100], using the PTM Scan Acetyl-Lysine Motif Kit (Cell Signaling Technology, 11663V1). The flow-through of the Acetyl-Lysine immunoprecipitation was kept for phosphorylated peptide enrichment. Eluted peptides were dried to completeness in a SpeedVac vacuum concentrator (Eppendorf) and subsequently resuspended in 0.1% TFA.

### Phosphopeptide enrichment

The peptide pool was acidified (pH<2) with 10% TFA and desalted using C18 cartridges (Sep-Pak Vac 1cc (50mg), Waters). Peptides were eluted with 3 × 150 µl 80% Acetonitrile (ACN) and 0.1% Formic Acid (FA), followed by freeze-drying. Enrichment of phosphopeptides was done by FeNTA beads (Thermo A-52283). Each of the 5 pools was enriched according to the manufacturer’s manual. The eluate was evaporated using a SpeedVac vacuum concentrator. Dried sample was dissolved in 50µL 01% TFA.

### Nano LC-MS/MS Analysis

The nano-HPLC system (Vanquish *Neo* UHPLC-System) was coupled to an Orbitrap Eclipse mass spectrometer, equipped with a FAIMS Pro Duo interface and a Nanospray Flex ion source (all parts Thermo Fisher Scientific). Peptides were loaded onto a trap column (Thermo Fisher Scientific, PepMap C18, 5 mm × 300 μm ID, 5 μm particles, 100 Å pore size) using 0.1% TFA as mobile phase. The trap column was switched in line with the analytical column (Thermo Fisher Scientific, PepMap C18, 75 μm ID × 50cm, 2 μm, 100 Å). The analytical column was connected to PepSep sprayer 1 (Bruker) equipped with a 10 μm ID fused silica electrospray emitter with an integrated liquid junction (Bruker, PN 1893527). Electrospray voltage was set to 2.4 kV. Peptides were eluted using a flow rate of 230 nl/min, starting with the mobile phases 98% A (0.1% formic acid in water) and 2% B (80% acetonitrile, 0.1% formic acid) and linearly increasing to 35% B over the next 180 min. This was followed by a steep gradient to 95%B in 5 min, stayed there for 5 min and ramped down in 2 min to the starting conditions of 98% A and 2% B for equilibration at 30°C. The Eclipse was operated in data dependent mode with a full scan (m/z range 350-1500, resolution 60,000, target value 4E5) at 3 different compensation voltages (CV-40, - 55, -70) followed by MS/MS scans of the most abundant ions at a cycle time of 1.0 s per CV. MS/MS spectra were acquired using an isolation width of 0.7 m/z, target value of 1E5 and intensity threshold of 2.5E4, maximum injection time of 120 ms, HCD with a collision energy of 36%, using the Orbitrap for detection, with a resolution of 50 k. For the detection of the TMT reporter ions, a fixed first mass of 110 m/z was set for the MS/MS scans. Precursor ions selected for fragmentation (including charge states 2-6) were excluded for 45 s. The Monoisotopic Precursor Selection (MIPS) mode was set to Peptide and the Exclude Isotopes feature was enabled.

### Peptide identification and quantification

For peptide identification, the RAW files were loaded into Proteome Discoverer (PD) (version 3.1.0.638, Thermo Scientific). All MS/MS spectra were searched using MSAmanda v 3.0.21.45 [101]. For peptide identification, precursor and fragment mass tolerance was set to ±10 ppm, with a maximum number of 2 missed cleavages, using tryptic enzyme specificity without proline restriction. RAW files were searched against the UniProt Physcomitrium patens database (v20210202, 61451 sequences), supplemented with common contaminants and protein tags. The iodoacetamide derivative on cysteine was specified as fixed modification, whereas oxidation on methionine, deamidation on asparagine and glutamine, acetylation on protein N-termini, carbamylation on lysine and peptide N-termini, glutamine to pyroglutamate conversion at peptide N-terminal glutamine and TMT modification at lysines and peptide N-termini were specified as variable modifications. For phospho-enriched samples, additionally phosphorylation on serine, threonine and tyrosine was considered as variable modification whereas acetylation on lysine was considered as variable modification in acetyl-enriched samples. Post-translational modification (PTM) sites within peptides were localized using ptmRS, which is based on the phosphoRS [102]. Searches of the input and enriched sample fractions (with additional modifications considered) were combined in a common Consensus Workflow in PD to ensure common protein grouping. Identifications were filtered to 1% FDR at the protein and PSM level using Percolator [103]. Furthermore, an Amanda Score cut-off of at least 150 was applied. Peptides were quantified based on reporter ion intensities extracted by the Reporter Ion Quantifier node implemented in PD. To distinguish regulated phospho-peptides that are altered due to regulation of the phosphorylation site from changes due to regulation of the underlying protein, PSMs were separated into those acquired from phospho-/acetyl-enriched samples and others acquired from the input before enrichments to capture changes on the proteome level. Therefore, proteins were quantified by summing unique and razor peptides detected and quantified in the 20 input fractions, whereas phospho- and acetyl-enriched samples were quantified on the peptide form level from the 5 enriched fractions respectively. Subsequently, phospho- /acetyl-peptide regulations determined in the enriched fractions were corrected by regulation of the corresponding protein in the input fractions to correct for changes observed due to regulation on the proteome level. For the subsequent analysis: contaminants, proteins with less than three detected peptides or PSMs, peptides where PTMs could not be localized to a specific amino acid, and peptides that could not be matched to a unique protein were excluded. Ensembl Gene IDs were retrieved by searching the UniProt IDs against the *Physcomitrium patens* V3.3 genome annotation. For volcano-plot visualization a Log10-P-value threshold of 1,3 were used with no Log2fold threshold.

### GO term annotation and enrichment analysis

For annotation, GO terms were retrieved by running protein IDs through UniProt ID Mapping. For GO enrichment analysis, the ShinyGO V0.80 (https://bioinformatics.sdstate.edu/go/) [104] tool was used. Default settings were used except for adjusting the maximum pathway size from 5000 to 2000 genes. A custom background was used consisting of all detected proteins from our proteome studies.

### Comparison to Published *P. patens* Acetylome and Proteome Studies

Comparison to published gametophore acetylome dataset [18] was done by searching acetylated proteins against unique proteins from the *rak1* hypoacetylome detected on our study. GO term enrichment was performed as described earlier. Comparison to published proteomic data on *P. patens* tissue type [50] was done by comparing the detected *rak1* proteome with published protonema and gametophore protein clusters. Overlap with the *rak1* proteome were standardized by filtering against a common protein background of 1800. To evaluate the statistical significance of the overlap, hypergeometric tests were performed. For each comparison, the observed number of overlapping proteins was compared to the expected number calculated as (n × D)/N, where n and D denote the sizes of the individual datasets and N represents the total number of proteins in the background. Hypergeometric p-values and a representation factor were computed to assess enrichment or depletion relative to random sampling from the common protein background.

### *In vitro* acetylation assay

Recombinant His6 tagged RAK1, NATDH protein was expressed in the pPICZA vector in *Pichia pastoris* BISY Mut^s^. Purification was done using a 1 ml prepacked IMAC column on an Äkta and eluted in 100mM Na-Pi, 100mM NaCl, 500mM Imidazole, pH 7.0. Recombinant *Arabidopsis* MKK1^DD^ (AT4G26070) was produced using a pOPNM vector in a BL12 codon plus E. coli strain with a MBP tag. MKK1^DD^ was purified using a GE Helthcare MBPTrap in 20mM Tris-HCl pH 7.4, 200mM NaCl, 1mM EDTA and 1mM DTT. Acetylation was done in tube using an acetyl transferase buffer (50mm 7,5ph tris, 10mM MgCl2, 10mM MnCl2, 1mM DTT, 50mM ATP, 0.1mM EGTA, 10% Glycerol) supplemented with 25µM [^14^C]-Acetyl-CoA (PerkinElmer), 200µM 19-mer H4 peptide and 400nM recombinant protein. After 1 hour, the H4 peptide was purified using SP sepharose resin (GE Heathcare) and the radioactivity was measured using a Hidex 300 sl scintillation counter.

### Structural modeling of RAK1

The structural model of RAK1 was generated by AlphaFold3 [105] under default settings using the amino acid sequence of RAK1 (Pp3c9_11360 / A0A2K1K2U5). PyMol software was used for visualization.

### List of Primers

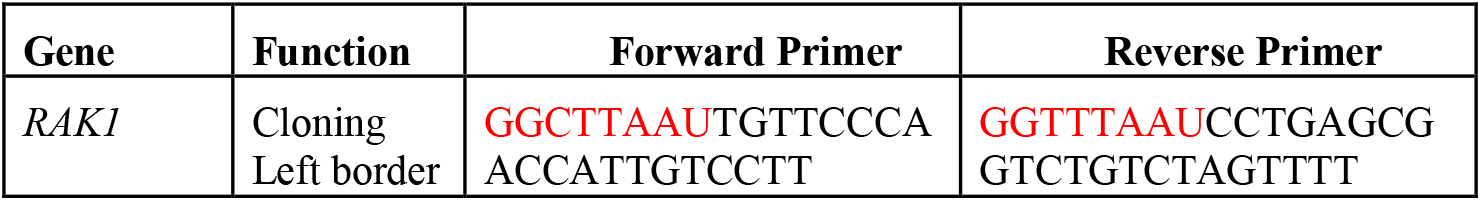
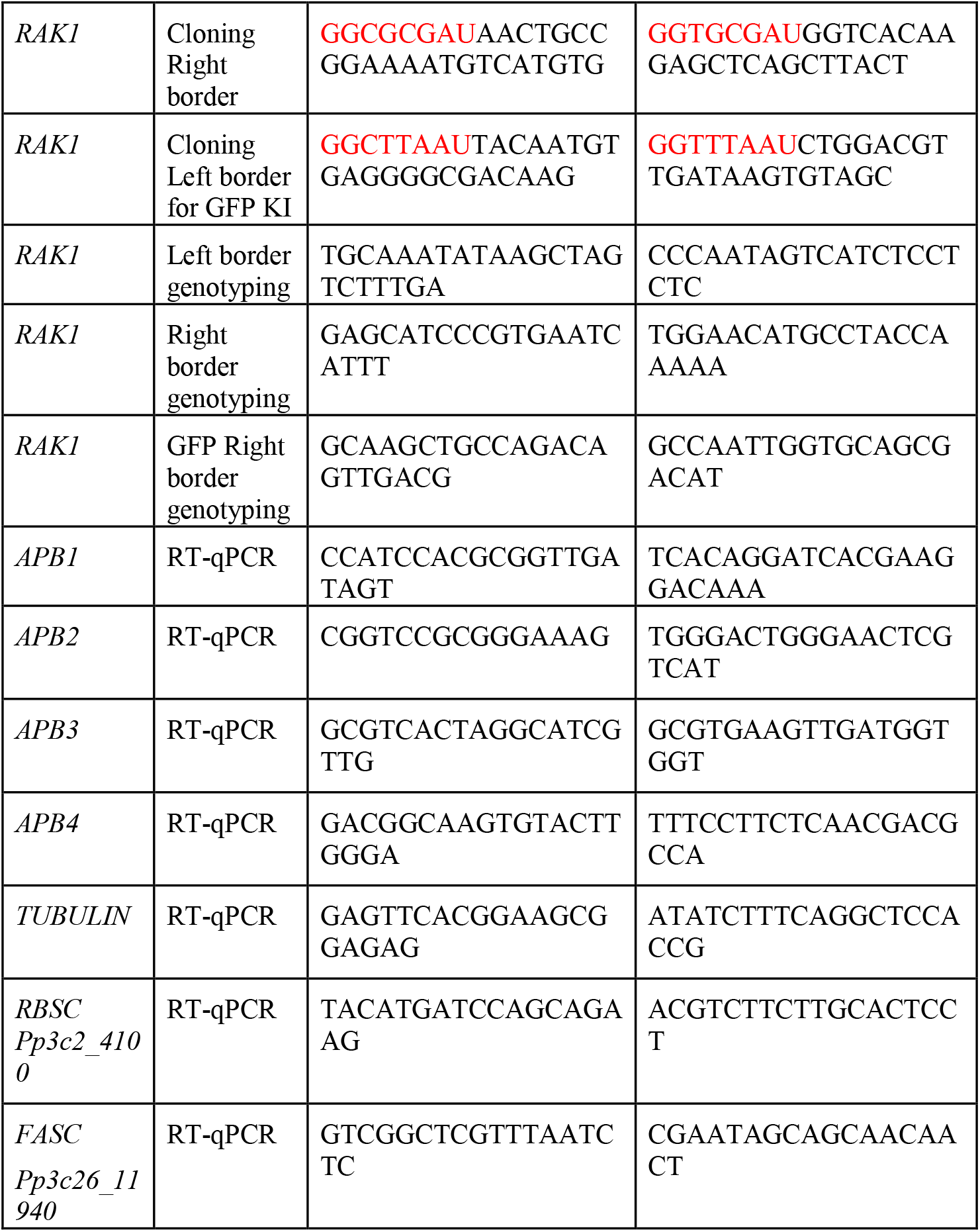

## Acknowledgments

We thank Jeppe Ansbøl, Emil Otto Kokholm Nielsen and Raquel Azevedo for valuable input and support during the bioinformatic analyses of the proteomic data and general handling of *Physcomitrium patens*. The authors also acknowledge John Mundy’s initial conceptualization of the project and Suksawad Vongvisuttikun for technical assistance.

## Funding

Danish Research Agency DFF1-1032-00249B (ER)

Novo Nordisk Fonden NNF190C0055222 (MP)

## Author contributions

Conceptualization: MP

Methodology: CDLH, TJA, JVK, MER, ZW, MS, GD, LZ, OS, MI

Investigation: CDLH, TJA, JVK, SS, MER, ZW, OS, MI, LAM

Visualization: CDLH, TJA, JVK, ZW

Supervision: MI, MH, YFD, LAM, ER, MP

Writing—original draft: CDLH, TJA, ER

Writing—review & editing: CDLH, TJA, LZ, MI, MH, LAM, ER, MP

## Data and materials availability

All data are available in the main text or the supplementary materials.

