## Supplemental Material for "A moss N-Acetyltransferase-MAPK protein controls 2D to 3D developmental transition via acetylation and phosphorylation changes"

Cloe de Luxán-Hernández *et al.*

**This PDF file includes:**

Figs. S1 to S4
Description of the Tables S1 to S20

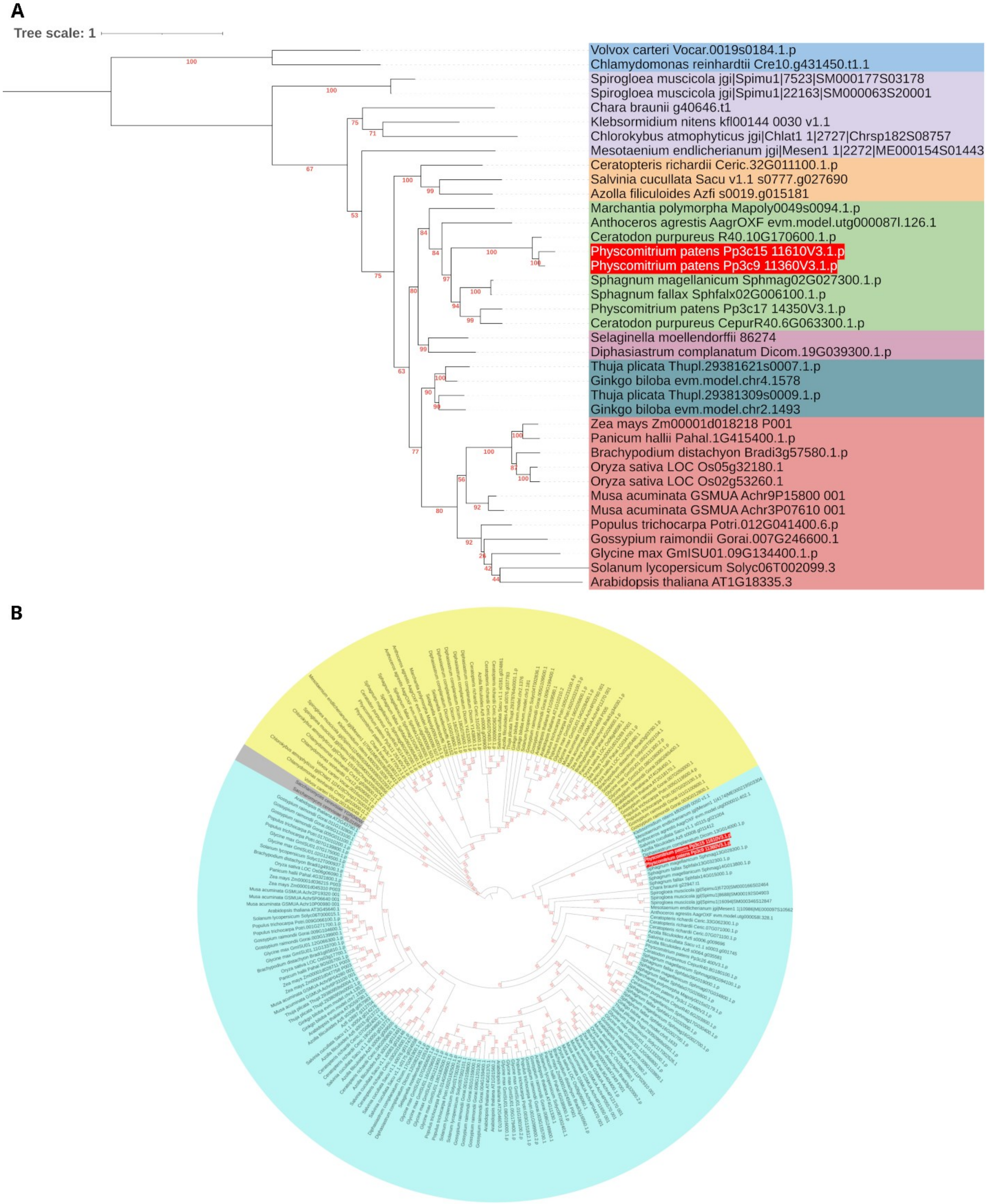


**Figure S1. Rossetta NATD-MAPK proteins are only present in mosses.**

(A) Phylogenetic tree based on multiple sequence alignment of NATDs found in OrthoFinder using land plant, green algae species and yeast as input. Complete protein sequence annotations are found in Table S1. Bootstrap values are shown at each node in red. Trees are drawn to scale with branch lengths measured in the number of substitutions per site. Clades are shown in different colours: Chlorophytes (blue), Charophytes (purple), Ferns (orange), Bryophytes (Green), Lycophytes (Pink), Gymnosperms (Teal) and Angiosperms (Red). RAK1 and RAK2 are highlighted in red. (B) Phylogenetic tree based on multiple sequence alignment of MAPKs found in OrthoFinder using land plant, green algae species and yeast as input. Complete protein sequence annotations are found in Table S2. Bootstrap values are shown at each node in red. Trees are drawn to scale with branch lengths measured in the number of substitutions per site. Clades are shown in different colours: Outgroup (Grey), Clade 1 (Yellow) and Clade 2 (Turquoise). RAK1 and RAK2 are highlighted in red.

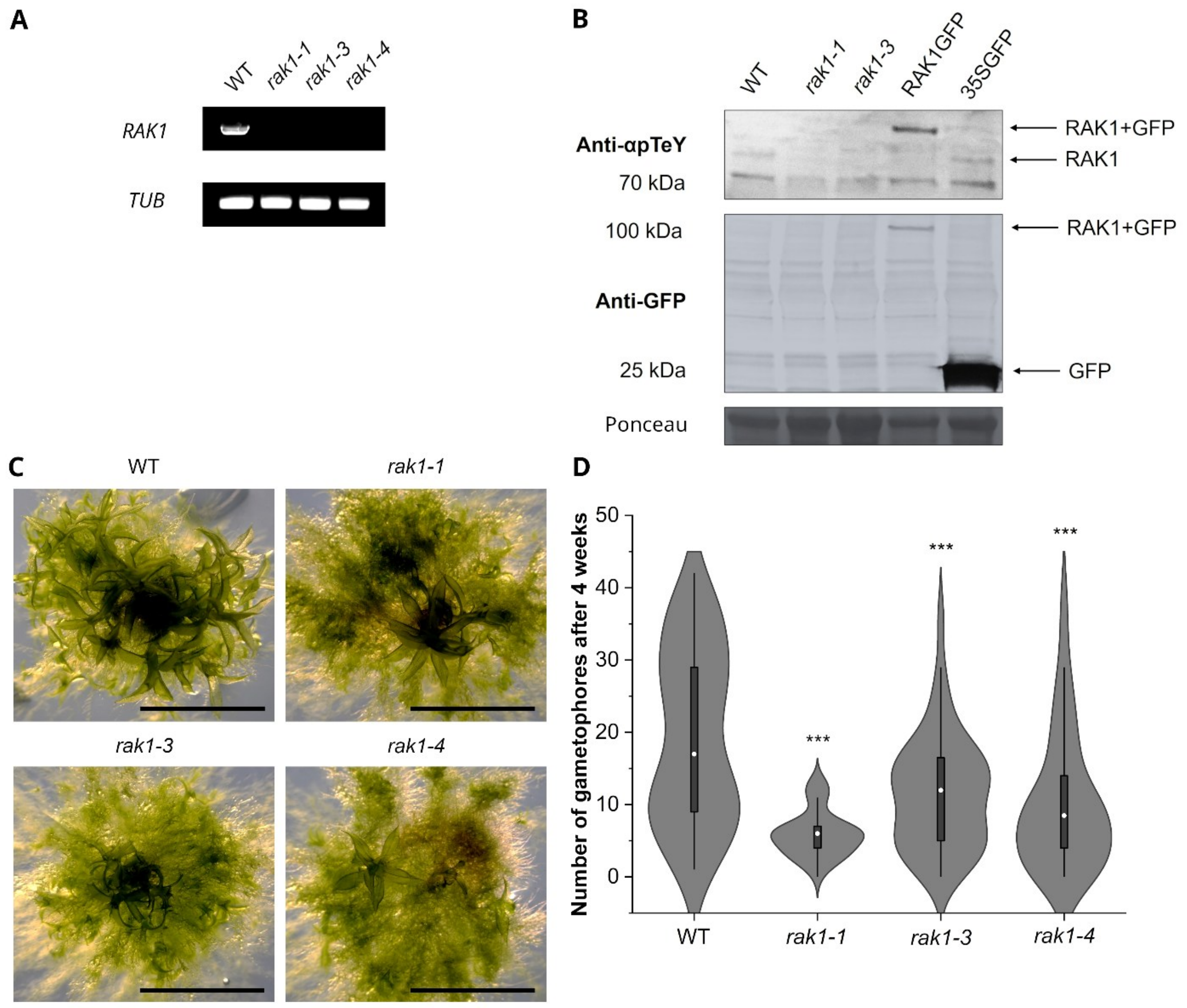


**Figure S2. *rak1* mutants develop less gametophores after 4 weeks with no apparent phenotypical defects.**

**(**A) RT-PCR showing that *rak1* generated mutants do not express the full transcript of *RAK1* (upper panel). *TUBULIN* expression was used as a loading control (lower panel). (B) Immunoblot analysis of RAK1 phosphorylation with α-pTEpY (Upper panel) and GFP-tagged proteins with α-GFP (Middle panel) in 2-week-old plants of the described genotypes. 35SGFP is used as a control for GFP detection. Lower panel shows ponceau staining of Rubisco, which is used as loading control. (C) Representative phenotypes of 4-week-old WT, *rak1-1*, *rak1-3* and *rak1-4* plants. Scale bar: 5 mm. (D) Quantification of the number of developed gametophores per plant: WT (N=48), *rak1-1* (N=49), *rak1-3* (N=52), and *rak1-4* (N=50). The box represents the interquartile range 25th to 75th. Median shown as a white circle. Whiskers represent 1.5 IQR. Statistical significance was calculated using a One-way ANOVA coupled to a Sidak test. Asterisks represent statistical difference against the WT: *rak1-1* (P value***: $4{,}39715\times10^{-12}$), *rak1-3* (P value***: $5{,}51497\times10^{-5}$), *rak1-4* (P value***: $1{,}56046\times10^{-6}$).

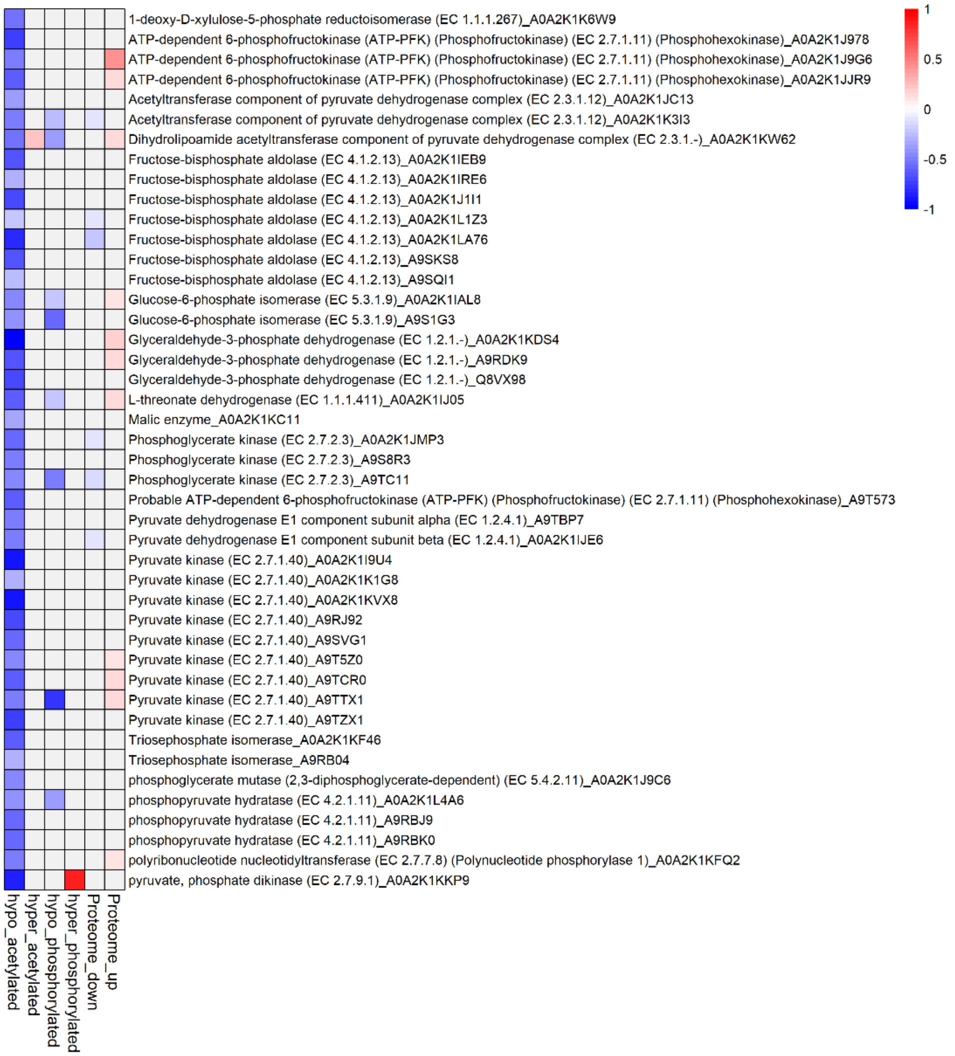


**Figure S3. Proteins involved in Acetyl-CoA synthesis are differentially hypoacetylated in *rak1*.**

Heat map showing differential $Log_2$Fold changes in *rak1-1* mutants in lysine acetylation, protein phosphorylation and protein abundance of proteins associated to the GO term “Pyruvate Metabolic Process”.

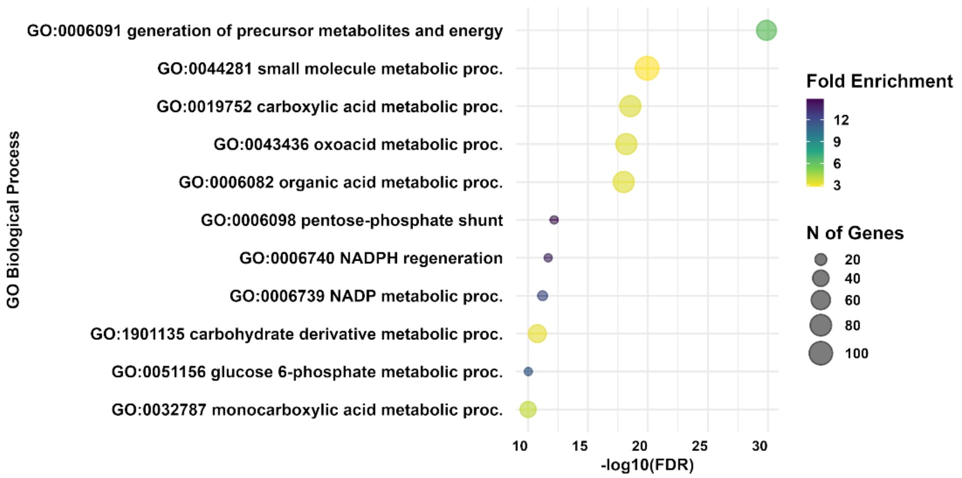


**Figure S4. Hypoacetylated proteins in *rak1* associated to gametophore-specific acetylation events are involved in metabolic processes.**

Representative top 11 significant Biological Process GO terms enriched in the overlapping proteins acetylated in gametophore tissue found by [18] and differentially hypoacetylated in *rak1-1*.

**Tables S1-20. (Excel file)**

**Table S1.** Protein sequences used for the Cobalt Alignment
**Table S2.** Protein sequences used to generate the Phylogenetic tree using the NATD domain of RAK1 as query
**Table S3.** Protein sequences used to generate the Phylogenetic tree using the MAPK domain of RAK1 as query
**Table S4.** Significant upregulated proteins in *rak1* vs WT
**Table S5.** Significant downregulated proteins in *rak1* vs WT
**Table S6.** Enriched GO terms of *rak1* vs WT upregulated proteins. GO term enrichment analysis using the ShinyGO V0.80 tool using setting described in the Materials and Methods section. A custom background was used consisting of all detected proteins from our proteome dataset
**Table S7.** Enriched GO terms of *rak1* vs WT downregulated proteins. GO term enrichment analysis using the ShinyGO V0.80 tool using setting described in the Materials and Methods section. A custom background was used consisting of all detected proteins from our proteome dataset
**Table S8.** Significant *rak1* vs WT hyperphosphorylated proteins with concatanated individual peptides
**Table S9.** Significant *rak1* vs WT hypophosphorylated proteins with concatanated individual peptides
**Table S10.** Enriched GO terms of *rak1* vs WT hyperphosphorylated proteome. GO term enrichment analysis using the ShinyGO V0.80 tool using setting described in the Materials and Methods section. A custom background was used consisting of all detected proteins from our proteome dataset
**Table S11.** Enriched GO terms of *rak1* vs WT hypophosphorylated proteome. GO term enrichment analysis using the ShinyGO V0.80 tool using setting described in the Materials and Methods section. A custom background was used consisting of all detected proteins from our proteome dataset
**Table S12.** Potential MAP Kinase targets. *rak1* vs WT hypophosphorylated proteins containing SP/TP sites
**Table S13.** Significant *rak1* vs WT hypoacetylated proteins with concataned peptides
**Table S14.** Significant *rak1* vs WT hyperacetylated proteins with concataned peptides
**Table S15.** Enriched GO terms of *rak1* vs WT hypoacetylated proteome. GO term enrichment analysis using the ShinyGO V0.80 tool using setting described in the Materials and Methods section. A custom background was used consisting of all detected proteins from our proteome dataset
**Table S16.** Proteins included in the *rak1* vs WT enriched GO-term "Pyruvate metabolic process" from the significant hypoacetylated data Table S13. Both proteome and phosphoproteome data is included
**Table S17.** Data used for heatmaps of *rak1* regulated Gametophore related proteins from Figure 5. For acetylated and phosphorylated proteins with several significant PTMs an average was made for the upregulated or downregulated parts respectively
**Table S18.** Comparison with *rak1* vs WT proteome and [50] protein tissue clusters. Overlapping proteins were found by first identifying a common background between both datasets
**Table S19.** Comparison with [18] data. Proteins shown are acetylated shared between the gametophore acetylome [18] and the *rak1* vs WT hypoacetylome (This study)

**Table S20.** Enriched GO terms of shared acetylated proteins between a published gametophore acetylome [18] and significant *rak1* vs WT hypoacetylated proteins proteome. GO term enrichment analysis using the ShinyGO V0.80 tool using setting described in the Materials and Methods section. A custom background was used consisting of all detected proteins from our proteome dataset
